# *STAT1* Gain-of-Function Variants Drive Altered T Cell Prevalence, Metabolism, and Heightened IL-6 Sensitivity

**DOI:** 10.1101/2021.11.10.468135

**Authors:** Saara Kaviany, Todd Bartkowiak, Daniel E. Dulek, Yasmin W. Khan, Madeline J. Hayes, Samuel Schaefer, Debolanle O. Dahunsi, James A. Connelly, Jonathan M. Irish, Jeffrey C. Rathmell

**Author notes:** Co-equal contribution (Saara Kaviany, Todd Bartkowiak).

## Abstract

Patients with Signal Transducer and Activator of Transcription 1 (*STAT1*) gain-of-function (GOF) pathogenic variants exhibit susceptibility to infections, autoimmunity, and cancer due to enhanced or prolonged STAT1 phosphorylation following cytokine stimulation. While interferons (IFNs) are canonical STAT1 activators, other cytokines that may also contribute to pathology in *STAT1* GOF patients have been less well defined. Here we analyzed the immune profiles and cytokine responses of two patients with heterozygous GOF mutations in the *STAT1* coiled-coil domain. A systems immunology approach revealed major changes in the T cell compartment and minor changes in the B cells, NK cells, and myeloid cells. Both patients with *STAT1* GOF differed from healthy individuals in the abundance and phenotype of effector memory, Th17, and Treg populations. *STAT1* GOF T cells displayed a pattern of increased activation and had elevated markers of glycolysis and lipid oxidation. Hypersensitivity of T cells to IL-6 was observed with intense, sustained STAT1 phosphorylation in memory T cell populations that exceeded that induced by IFNs. Together, these results show a role for STAT1 in T cell metabolism and suggest that IL-6 may play a critical role to promote T cell memory formation and activation in patients with *STAT1* GOF.

## Introduction

A class of recently discovered inborn errors of immunity (IEIs) involve abnormal cellular signaling through mutations in members of the Signal Transducer and Activator of Transcription (STAT) family. STAT proteins are latent transcription factors that require cytokine stimulation to be recruited to receptors via their SH2 domains. Subsequent tyrosine phosphorylation by Janus kinases (JAKs) leads to formation of multimers and translocation into the nucleus, where the STAT complexes bind specific DNA sequences and regulate gene transcription. JAK/STAT signaling pathways are activated by a wide range of cytokines, with STAT1 phosphorylation occurring following cell stimulation with Interferons α, β, γ, λ, IL-27, and IL-6[1–4]. *STAT1* gain-of-function (GOF) mutations are recognized as IEIs that present with a wide range of clinical phenotypes [5]. Characterization of 274 patients with pathogenic *STAT1* GOF mutations identified significant infectious complications within this patient cohort including bacterial, viral, chronic mucocutaneous candidiasis (CMC) and invasive fungal infections [6]. Patients with *STAT1* GOF mutations may also have autoimmune manifestations, including hypothyroidism, type 1 diabetes, hematologic cytopenias, cancers, and systemic lupus erythematosus (SLE) [5, 6]. Invasive infections, cerebral aneurysms and cancer were the strongest predictors in poor outcome in affected patients [6]. Importantly, neither genotype-phenotype correlation nor the risk factors for poor outcomes in *STAT1* GOF patients have been described to date. Moreover, the mechanisms that drive these diverse phenotypes remain uncertain.

Pathogenic *STAT1* GOF mutations lead to increased STAT1 phosphorylation [3, 5, 7] and increased downstream phosphorylated STAT1 (p-STAT1)-driven target gene transcription. Basal STAT1 phosphorylation may, however, be modest and increased activity can require cytokine stimulation. Processes that lead to STAT1 hyperphosphorylation in response to cytokines include impairment of dephosphorylation, increased STAT1 protein levels, and premature nuclear import of p-STAT1 [8]. Increased STAT1 signaling may affect development of specific T cell subsets and low levels of circulating Interleukin-17A-producing T cells are identified in the large majority (82%) of patients and contributes to the predisposition to fungal infection in these families [6]. However, the predisposition to viral and intracellular infections which depend upon strong Type I and Type II interferon responses remains a paradox without clear explanation.

Here we applied a systems immunology approach to deeply characterize signaling, metabolism, activation state, and differentiation state of the peripheral blood immune cells altered by STAT1 variants. Given the limited characterization of the immune milieu in these patients, we used high-dimensional mass cytometry to comprehensively characterize the lymphoid and myeloid compartments in two patients with *STAT1* GOF mutations. Using this approach, we were able to track major mononuclear cell types frequencies and activation states, including phosphorylation and metabolism [9–14], and cytokine responses in two *STAT1* GOF patients. These studies demonstrate the effect of pSTAT1 perturbations on a global cellular level and identified the impact of *STAT1* GOF in specific cell subsets. While all immune cell types were affected, shifts in T cell subsets and activity were notable, with decreased effector memory and Treg and altered markers of T cell metabolism that resemble early activated T cells. These studies confirmed and provided new detail on altered T cell populations in patients with these variants and identified a highly increased responsiveness to IL-6 that may have important clinical implication.

## Results

### Patient descriptions

We studied two patients with heterozygous *STAT1* GOF mutations and distinct clinical histories to further define potential immunopathogenic mechanisms of *STAT1* GOF-associated disease **(Supplementary Table 1)**. Patient 1 (P1) presented as a previously healthy male at 9-months of age with persistent fever. Laboratory and clinical evaluation demonstrated pancytopenia, a significant transaminitis, elevated ferritin, hepatosplenomegaly, and coagulopathy to meet diagnostic criteria for hemophagocytic lymphohistiocytosis (HLH) syndrome. Evaluation of infectious triggers of HLH identified disseminated histoplasmosis through detection of histoplasma antigen in peripheral blood. He was successfully treated with dexamethasone to control his elevated immune response, as well as liposomal Amphotericin B with transition to itraconazole to successfully treat the disseminated histoplasmosis. Clinical immunologic evaluation at the time of initial presentation demonstrated decreased NK function by clinical chromium release assay, but normal CD107a degranulation. Clinical B-cell phenotyping revealed low frequencies of non-switched memory B-cells, switched memory B-cells and total memory B-cells. T cell functional testing demonstrated moderately decreased CD3+ T cell proliferation in response to tetanus toxin, and normal mitogen stimulation to phytohemagglutinin (PHA) and pokeweed mitogen (PWM). He demonstrated a modestly decreased CD45+ total lymphocyte proliferation to Candida, however his CD3+ T cell response was normal, overall suggesting a normal response. A primary immunodeficiency next generation sequencing (NGS) panel identified a *de novo* pathogenic variant in *STAT1* (c.800C>T; p.Ala267Val) previously identified in patients with CMC to result in GOF [15–19]. This missense mutation in the coiled-coiled domain has previously been shown to lead to enhanced phosphorylation in response to IFN (α and γ) and to impair dephosphorylation of p-STAT1 protein, leading to the described GOF [20, 21]. PBMC samples were collected 6 months after discontinuing immune suppression and when the patient had controlled his histoplasma infection in attempts to minimize the impact of recent dexamethasone, however the patient remained lymphopenic at the time of collection.

Patient 2 (P2) is male and presented at 8 years of age with an episode of HSV dermatitis of the eyelid followed by aseptic meningitis. Although VZV was not directly detected in his cerebrospinal fluid (CSF) via PCR, his VZV CSF IgG index was elevated, suggestive of VZV meningitis. Immune evaluations demonstrated low frequency and absolute number of NK cells with diminished function by chromium release assay, yet normal CD107a expression. T cell and B cell populations were quantitatively within normal range, including memory and naïve subsets. He demonstrated normal immunoglobulin levels, with appropriate post-vaccine titers. A primary immunodeficiency NGS panel revealed a pathogenic variant in *STAT1*, c.866A>G (p.Tyr289Cys) in the coiled-coiled domain. Familial testing was not performed. This variant has been reported to increase STAT1 phosphorylation and has also been reported in individuals with CMC [3, 6]. PBMCs from P2 were collected at a baseline several years after treatment for the episodes of HSV dermatitis and VZV meningitis and while not experiencing detectable immune or infectious complications.

### STAT1 GOF Shifts T cell Subsets and Activation States

We performed detailed immunophenotyping with high dimensional cytometry to determine how *STAT1* GOF influenced immune cell populations in these two patients (gating schema in **Supplemental Figure 1)**. T cells from five healthy donors, P1, and P2 were analyzed with mass cytometry using a panel of antibodies designed to define T-cell subsets, activation state, and metabolism (**Supplemental Table 2)**. Despite different clinical phenotypes, dimensionality reduction using tSNE plots in P1 and P2 demonstrated several similarities when compared to healthy individuals (**Figure 1a**). For each patient with a *STAT1* GOF mutation, the T cell populations in the tSNE plots demonstrated a notable alteration in T cell memory populations compared to healthy donors. Both patients had reduced effector memory (Tem) CD4+ and CD8+ T cell populations, yet the frequencies of naïve (Tn) and central memory (Tcm) did not appear to follow the same pattern **(Figure 1b)**. In particular, the frequency of CD8+ Tn was elevated in P1 (72%) and P2 (88%) compared to the median frequency of CD8+ Tn in healthy donors (57%, IQR 23%) **(Figure 1b)**. P1 had 6% CD8+ Tem, P2 had 8% compared to median of 13% CD8+ Tem in healthy donors. The CD4+ compartment demonstrated similar patterns in the frequency of memory populations. In P1, the CD4+ pool consisted of 74% Tn and 9% Tem, with similar findings in P2 with 71% CD4+ Tn and 5% Tem. The healthy donors, however, showed a trend for fewer CD4+ Tn (54%) and more Tem (16%).

**Figure 1.**
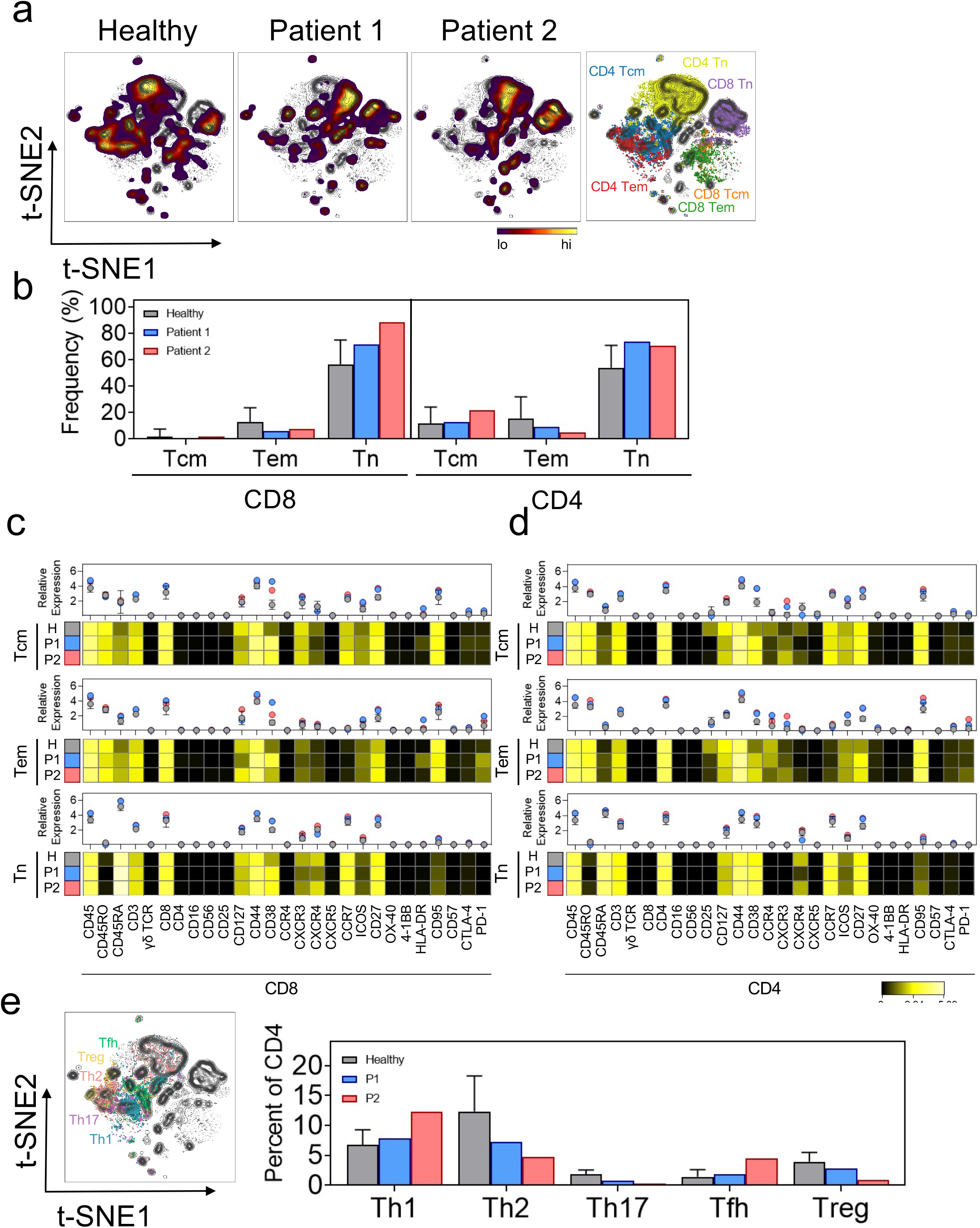
*STAT1* GOF Patients Demonstrate Comparable Alterations in Memory, Helper, and Activated T cell subsets. **a**. Representative t-SNE plots depicting the density of CD45+CD3+ cells (heat) from each individual healthy donor (far left), patient 1 (middle left), or patient 2 (middle right). Density contours indicate the cells from each patient overlaid onto a contour plot of cells pooled from each sample. T cell memory populations are indicated in manually gated populations overlaid onto the t-SNE plot on the far right. **b-d. (b)** Bar charts demonstrating the frequency of CD8+ (left) and CD4+ (right) central memory (Tcm), effector memory (Tem) and naïve T cell (Tn) populations in five healthy controls and two patients with *STAT1* GOF mutations. Heat maps indicating the arcsinh transformed median values of 27 phenotypic markers expressed in CD8+ Tcm, Tem, or Tn populations (**c**) or CD4+ T cells (**d**) manually gated from healthy donor PMBC (grey), P1 (blue) or P2 (red). Graphs above each heatmap indicate the arcsinh fold change expression of each marker relative to the minimum median mass intensity. **e**. Representative t-SNE plot (left) overlaying manually gated helper T cell populations onto pooled CD45+CD3+ cells as in **a**. Bar graphs (right) indicate the frequency of each manually gated T helper subset within the CD4+ T cell pool. All values represent the median statistic. Errors bars indicate the interquartile range.

Not only did T cell memory frequencies appear altered in the patients with *STAT1 GOF*, but T cells appeared more activated in these patients. Both CD4+ and CD8+ Tcm and Tem from P1 and P2 had elevated levels of a wide range of activation markers, including CD44, CD27, and CD38 **(Figure 1c, d)**. Across Tcm, Tem, and Tn CD8+ T-cells, activation markers CD44, CD38, and CD27 were more highly expressed in the patients with *STAT1* GOF compared to healthy individuals. Specifically, CD8+ Tcm from both patients expressed higher levels of CD44 (P1: 1.2-fold; P2: 1.1-fold), CD38 (P1: 3-fold; P2: 2-fold), ICOS (P1: 2-fold; P2: 2-fold) and CD27 (P1: 1.5-fold; P2: 1.6-fold) compared to healthy individual populations **(Figure 1c)**. Similar trends were identified in CD8+ Tem and Tn populations as well as Tcm, Tem and Tn subsets in the CD4+ T-cell compartment **(Figure 1d)**. Interestingly, PD-1 was more abundantly expressed on Tem in the patients with *STAT1* GOF compared to healthy individuals. Specifically, PD-1 was upregulated in the CD8+ Tem cells for both *STAT1* GOF patients (P1: 3 fold, and P2: 2 fold) and was elevated in CD4+ Tem population of P2 (1.8 fold). These data show a decreased abundance of Tem but suggested increased activation state or chronic stimulation of the remaining Tem caused by *STAT1* GOF.

The impact of *STAT1* GOF mutations on T-helper differentiation into functional subsets was next investigated. Using surface markers characteristic of CD4+ subsets, effector (Th1, Th2, Th17, Tfh) and regulatory (Treg) populations in P1 and P2 were compared to healthy donors (**Figure 1e**, **Supplementary Figure 1a**). While the frequencies of CXCR3+ Th1-like T cells were variable and not significantly different from healthy individuals, CCR4+ Th2-like cells appeared to be diminished in each patient. Th17 cells were also less frequent in *STAT1* GOF patients, consistent with what has been previously described in *STAT1* GOF mutations [6]. Notably, the circulating CXCR5+PD-1+ T follicular helper cells (Tfh) population demonstrated the opposite pattern, and positive Tfh cells were elevated in P1 and P2 compared to healthy individuals. Lastly, the frequency of regulatory T-cells (Tregs) was diminished in both P1 and P2, accounting for less than 1% of the CD4+ T cell populations in P2 and less than 3% in P1, compared to 4% in healthy donors.

### STAT1 *GOF and B cell, NK, and Myeloid Immunophenotyping*

Given the alterations of T cell populations caused by *STAT1* GOF mutations, we next examined other immune cell populations in these patients. The T cell panel of antibodies for mass cytometry also included the NK cell markers CD16 and CD56 and these data were also examined to assess potential changes in NK cell populations. No consistent difference in abundance of CD56^bright^ and CD56^dim^ NK cells was observed between the two *STAT1* GOF patients compared to healthy donor (**Supplementary Figure 1a, 2**). In both cases, however, NK cells in the *STAT1* GOF patients had elevated expression of CD127, CD95, and HLA-DR to suggest activated states. A separate panel of antibodies focused on B cell surface markers was used to identify B cell populations by mass cytometry (**Supplemental Table 2)**. Dimensionality reduction using tSNE plots showed distinct patterns in the frequencies of B cell subsets in patients with *STAT1* GOF (**Figure 2a**). Identification of each population showed that while the frequencies of most subsets were unchanged, class switched memory B cells were greatly reduced to only 3% in P1 and P2, compared to 16% in healthy individuals (**Figure 2b, Supplementary Figure 1b**). Beyond decreased abundance of this population, few individual markers were consistently altered within a given population except for CD38, which was increased in the class switched B cells that were present in patients with *STAT1* GOF (**Figure 2c**). Finally, a myeloid cell-focused panel of antibodies was used to examine these innate cell populations (**Supplemental Table 2**). Myeloid cell populations were less consistent than lymphoid populations, although there appeared to be a trend towards increased frequencies of inflammatory monocytes (i-mono) and classical dendritic cell (cDC) populations with *STAT1* GOF (**Figure 3a, b, Supplementary Figure 1c**). Expression of S100A9 and HLA-DR were widely elevated to suggest myeloid cells were also broadly activated (**Figure 3c**).

**Figure 2.**
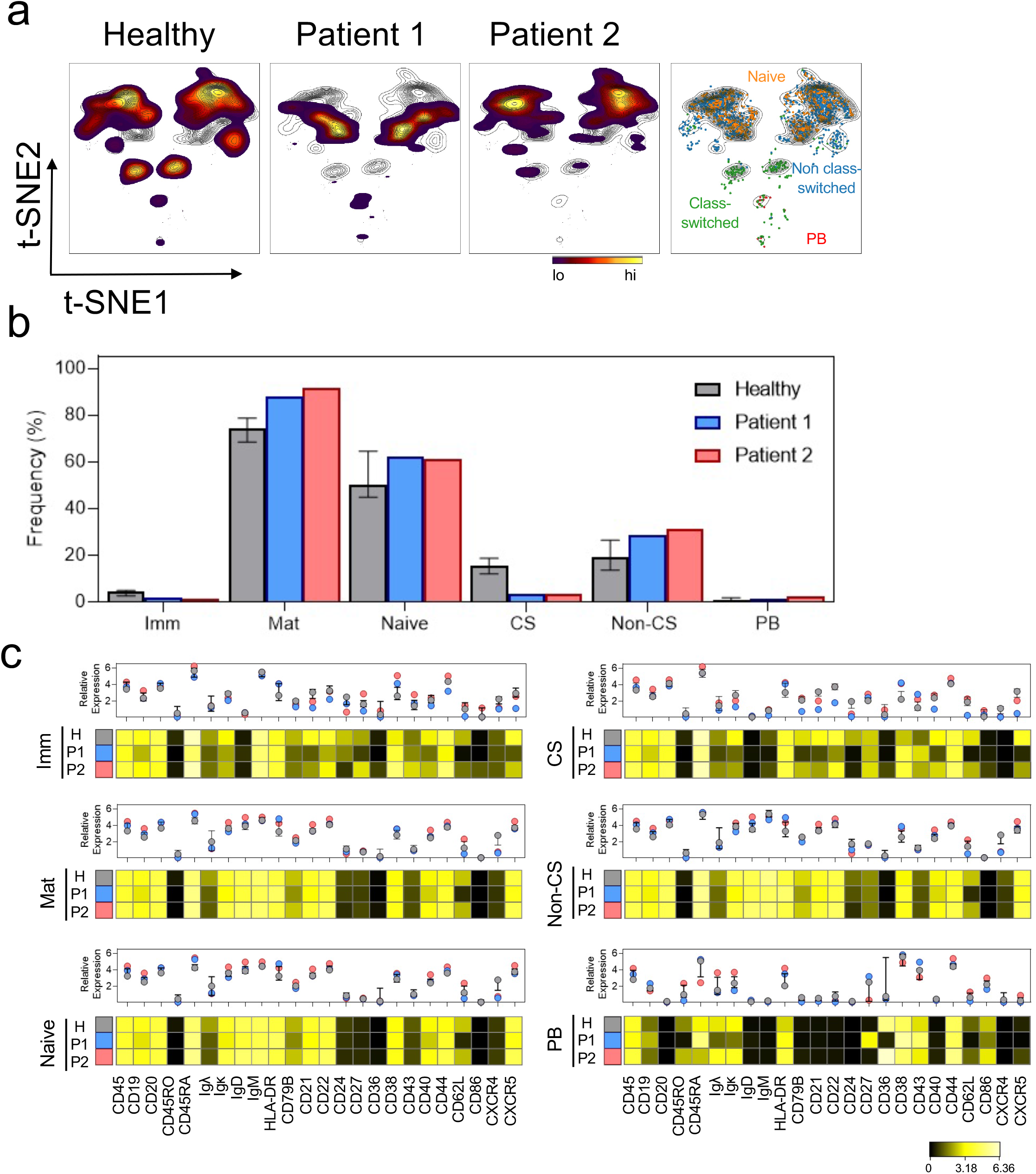
Deficit of Class Switched B cells in Patients with *STAT1* GOF mutations. **a**. Representative t-SNE plots depicting the density of CD45+CD19+CD20+ cells (heat) from each individual healthy donor (far left), P1 (middle left), or P2 (middle right). Density contours indicate the individual patient’s cells overlaid onto a contour plot of cells pooled from each sample. B cell populations are indicated in manually gated subsets overlaid onto the t-SNE plot on the far right. **b**. Bar charts demonstrating the frequency of manually gated B cell populations in five healthy controls and two patients with *STAT1* GOF mutations. **c**. Heat maps indicating the arcsinh transformed median values of 24 phenotypic markers expressed in manually gated populations of immature B cells (imm), mature B cells (Mat), naïve B cells (Naïve), class-switched B cells (CS), non-class switched B cells (non-CS) and plasmablasts (PB). Graphs above each heatmap indicate the arcsinh fold change expression of each marker relative to the minimum median mass intensity. All statistics represent the median value. Errors bars indicate the interquartile range.

**Figure 3.**
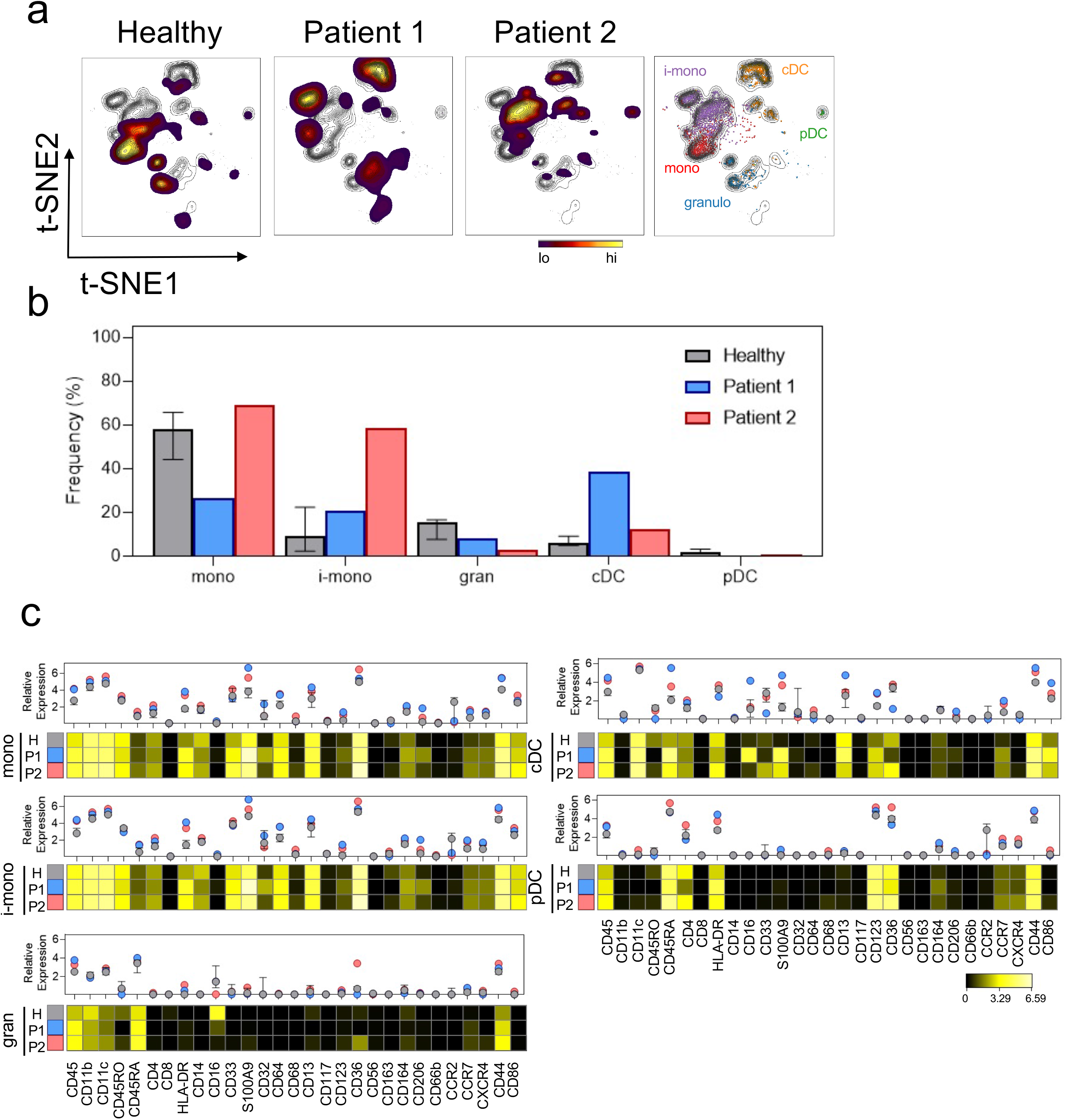
Myeloid Populations in Patients with *STAT1* GOF Mutations have Increased Immature Monocytes and Classic Dendritic Cells. **a**. Representative t-SNE plots depicting the density of CD45+CD3-CD19-CD56-cells (heat) from each individual healthy donor (far left), P1 (middle left), or P2 (middle right). Density contours indicate the individual patient’s cells overlaid onto a contour plot of cells pooled from each sample. Myeloid populations are indicated in manually gated subsets overlaid onto the tSNE plot on the far right. **b**. Bar charts demonstrating the median frequency of manually gated B cell populations in five healthy controls and two patients with STAT1 GOF mutations. **c**. Heat maps indicating the arcsinh transformed median values of 29 phenotypic markers expressed in manually gated populations of monocytes (mono), inflammatory monocytes (i-mono), granulocytes (gran), classical dendritic cells (cDC) and plasmacytoid dendritic cells (pDC). Graphs above each heatmap indicate the median arcsinh fold change expression of each marker relative to the minimum median mass intensity. All statistics represent the median value. Errors bars indicate the interquartile range.

### Increased Markers of Active Cell Metabolism in STAT1 GOF

T cell metabolism increases upon stimulation to support bioenergetics, biosynthesis, ad cell signaling[22]. Early after activation both glucose and mitochondrial metabolism increase, with glucose metabolism decreasing and mitochondrial lipid oxidation becoming dominant as T cells transition to memory phenotypes [9]. To assess how *STAT1* GOF affected these metabolic processes, expression of the Glucose transporter 1 (Glut1) and carnitine palmitoyl transferase 1A (CPT1a) was measured by mass cytometry. Interestingly, both patients with *STAT1* GOF mutations expressed higher glucose transporter 1 (Glut1) and carnitine palmitoyl transferase 1A (CPT1a) across different cell subtypes compared to healthy individuals (**Figure 4a, b**). Monocytes and dendritic cells expressed the highest levels of both Glut1 and CPT1 that corresponded to the highest levels of p-STAT1 and p-STAT3. Within the CD4+ and CD8+ memory populations, Tcm expressed the highest levels of Glut1, and Tem and Temra populations expressed the highest levels of CPT1a compared to healthy individuals (**Figure 4c**). These data may suggest early activation states or a failure of memory T cells to downregulate Glut1 after prolonged stimulation. Regulatory T-cells are less reliant on glycolysis with low Glut1 expression [23], but Tregs from patients with *STAT1* GOF, in particular P2, had elevated expression of Glut1 that was comparable to effector T cell populations and may indicate reduced suppressive capacity [24](**Figure 4d)**. In the two patients with *STAT1* GOF mutation, this overall pattern of increased expression of Glut1 and CPT1a compared to healthy donor suggests an activated T cell state and a key role of STAT1 to mediate immune metabolic phenotypes.

**Figure 4.**
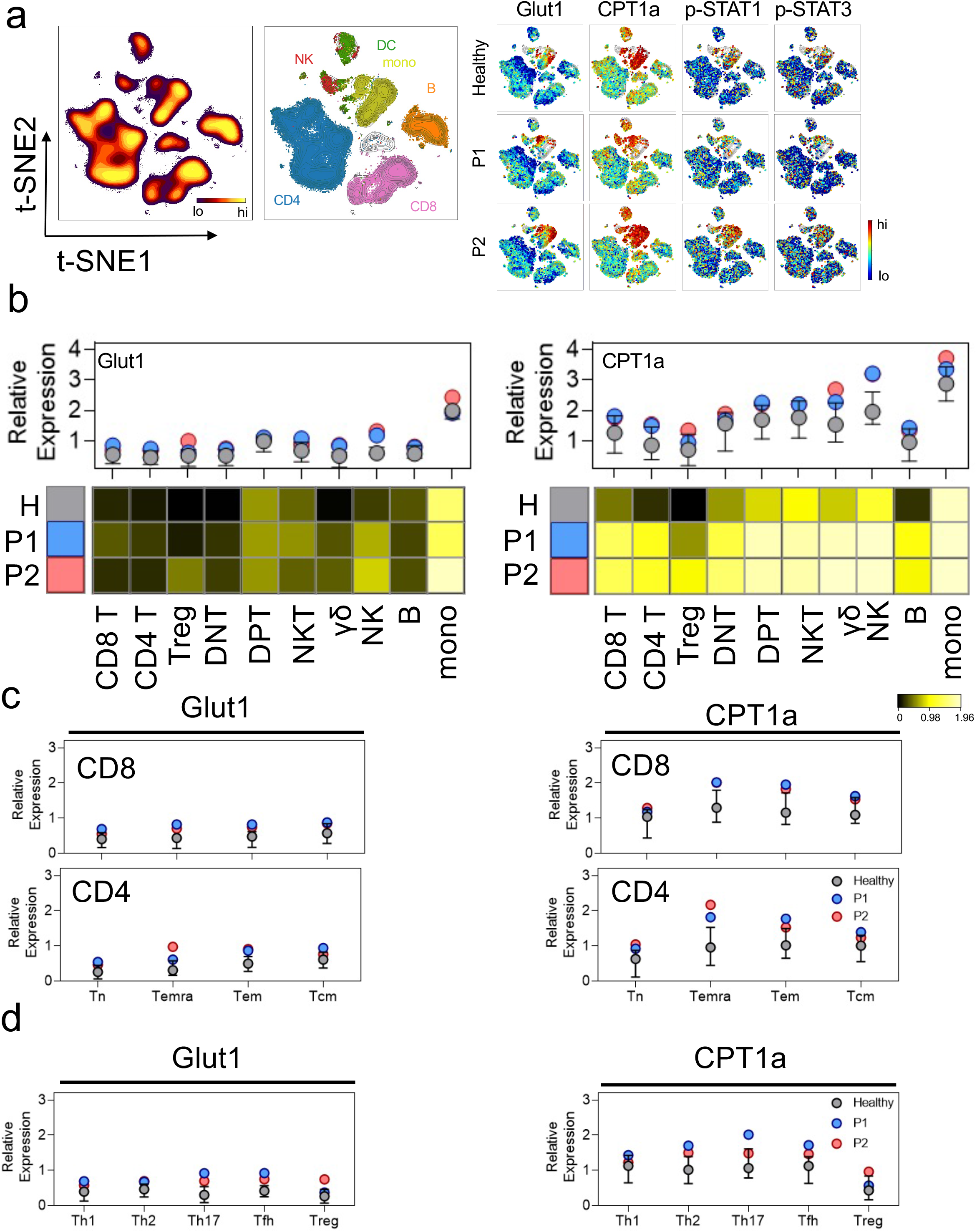
Increased Expression of Cell Metabolic Markers in Patients with *STAT1* GOF Mutations. **a**. Representative t-SNE plot depicting the density of CD45+ cells (heat) pooled from one healthy donor and each patient (left). Manually gated populations were overlaid onto t-SNE contours indicating each immune subset’s position on the t-SNE plot. (Right) Heatmaps of indicated expression values of indicated markers on the t-SNE plot. **b**. Heatmaps indicating the arcsinh transformed median expression values of Glut1 (left) and CPT1a (right) on indicated immune populations in healthy donors and STAT1 GOF patients. Graphs above each heatmap indicate the median arcsinh fold change expression of each marker relative to the minimum median mass intensity across all immune subsets. **c**. Quantification of the median arcsinh transformed intensities of Glut1 (left) and CPT1a (right) in manually gated CD8+ (top) and CD4+ (bottom) naïve and memory populations. **d**. Quantification of the median arcsinh transformed intensity values of Glut1 (left) and CPT1a (right) in manually gated T helper subsets as indicated. All statistics represent the median value. Errors bars indicate the interquartile range.

### Stimulation with IFNγ and IL-6 demonstrates Differential Phosphorylation of STAT1 GOF

Given the broad pattern of activated immune cells, we next explored the phosphorylation of STAT proteins within *STAT1* GOF peripheral blood mononuclear cells (PBMCs) to cytokine stimulation. To assess the impact of these two *STAT1* GOF mutations on immune function, we first compared the levels of basal STAT1 phosphorylation in lymphoid and myeloid subsets (**Figure 4a and Supplemental Figure 3**). Both patients with *STAT1* GOF mutations exhibited increased basal phosphorylated STAT1 (p-STAT1) compared to healthy donor individuals in T, B, NK, and myeloid cells, with T cells and NK cells appearing most strongly affected (**Supplementary Figure 3)**. T cells from P1 exhibited 5-fold higher basal p-STAT1 compared to the median p-STAT1 levels in healthy donors (0.1, IQR= 0.3), whereas T cells from P2 exhibited a 7-fold increase in basal p-STAT1 compared to the median phosphorylation of healthy donor T cells. Basal STAT1 signaling did not appear greatly impacted in the B cell compartment in either patient, as B cells had only a modest increased in p-STAT1 signaling of 1.4-fold (P1) and 1.9-fold (P2) increase over the median p-STAT1 compared to healthy controls. NK cells from P1 exhibited 2.9-fold increased STAT1 phosphorylation and NK cells from P2 displayed 4-fold increased phosphorylation compared to the median of healthy donor controls (0.2, IQR= 0.4). Myeloid cells had the greatest basal STAT1 phosphorylation (median=0.4; IQR=0.6). Like the lymphoid populations, myeloid cells expressed 2-fold greater p-STAT in P1 and 3-fold higher in P2. Interestingly, basal p-STAT3 and p-STAT5 were also increased in both patient samples across these immune cell types, indicating broadly increased STAT signaling and immune activity in the patients with *STAT1* GOF mutations that may be a consequence of altered cytokine profiles and signaling. These data show that *STAT1* GOF mutations can have direct and indirect impact on other STAT signaling pathways.

We next tested the STAT activity of different immune cell populations in response to cytokine stimulation. PBMCs were left unstimulated or treated for 15 minutes with IFNa and IFNγ, cytokines that classically signal directly through STAT1, interleukin-6 (IL-6) which classically signals through STAT3 but can also activate STAT1, or hydrogen peroxide to broadly inhibit phosphatases and act as a positive control. Cells were analyzed by mass cytometry using phospho-specific antibodies to measure the phosphorylation of STAT1 and STAT3 in distinct cell populations (**Figure 5a, b, c, Supplementary Table 2**). P1 generally failed to induce extensive phosphorylation of STAT1 or STAT3 or exhibited modest response to all stimuli, including the positive control hydrogen peroxide. It was possible the 15-minute stimulation time point was too brief and P1 would have delayed STAT phosphorylation, CyTOF was insufficiently sensitive, or compensatory mechanisms may dampen all STAT signaling in P1.

**Figure 5.**
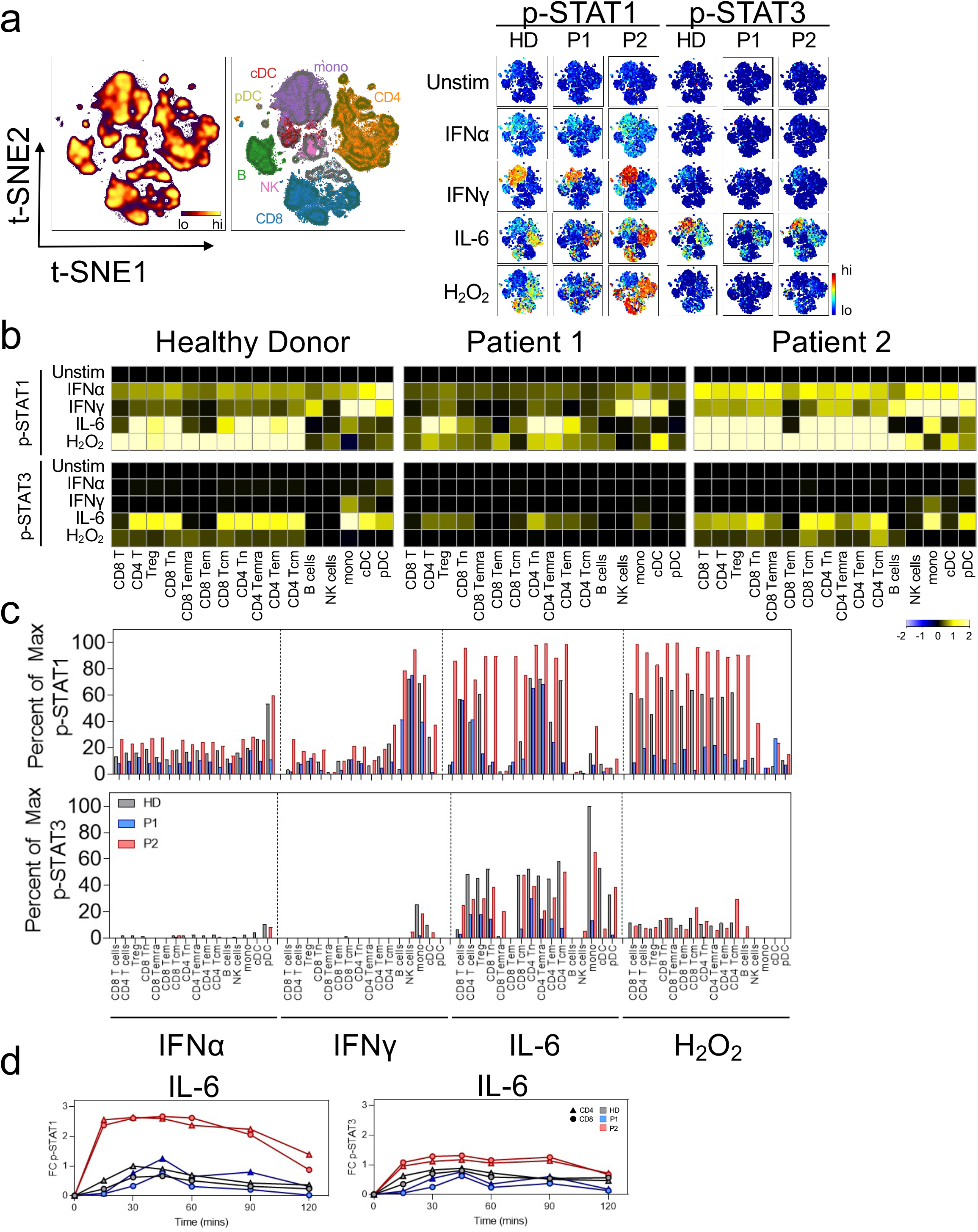
Stimulation with IFN-γ and IL-6 Demonstrates Differential Phosphorylation of STAT in *STAT1* GOF Patients. **a**. Representative t-SNE plot indicating the density of each CD45+ immune population in the graph (left). Manually gated populations were overlaid onto t-SNE contours indicating each immune subset’s location along the t-SNE axes. Heatmaps on the t-SNE axes indicating the expression of p-STAT1 or p-STAT3 under each given stimulation condition. **b**. Heatmaps indicate the median arcsinh transformed expression values of p-STAT1 (top) or p-STAT3 (bottom) within each indicated manually gated immune population exposed to the indicated cytokine for 15 minutes. Each value is normalized to each individual’s unstimulated condition. **c**. Quantification of the percent of the maximum possible p-STAT1 signal (top) or p-STAT3 signal (bottom) achieved across all manually gated immune subsets and stimulation conditions as indicated. **d**. Time course of p-STAT1 (right) or p-STAT3 signaling in response to IL-6 stimulation. All phospho-protein values were arcsinh transformed and normalized to each individual’s baseline (timepoint 0) median intensity values. Triangles indicate manually gated CD4+ T cells. Circles indicate manually gated CD8+ T cells.

Given the possibility that immune cells in *STAT1* GOF mutations may show altered STAT1 phosphorylation kinetics, we next assessed the temporal regulation of STAT phosphorylation conferred by these STAT1 variants. STAT phosphorylation in P1 was tested at multiple time points using fluorescent flow cytometry following treatment with IFNγ, IL-6, or IL-10 (**Supplemental Figure 4**). P1 did induce STAT1 and STAT3 phosphorylation, but these were similar in level to those of heathy individuals. In contrast to P1, IFNα and IFNγ led to robust STAT1 phosphorylation without increased p-STAT3 in healthy donor and P2 **(Figure 5a, b, c)**. This was less clear for pDCs and monocytes/cDCs which showed modest IFN-induced p-STAT3 to IFNα and IFNγ respectively. P2 had the greatest IFNα and IFNγ-induced phosphorylation of STAT1 across all T cell types and subsets. NK cells from P2 also showed heightened IFNγ-induced STAT1 phosphorylation.

IL-6 can also activate STAT signaling. While IL-6 canonically activates STAT3 and this response was similar or less effective in P1 and P2, IL-6 induced a strong STAT1 phosphorylation in CD4+ and CD8+ T cell subsets in P2. The healthy control patients demonstrated an expected increase in pSTAT1 following IL-6 stimulation. This response was less prominent in CD4+ effector memory (Tem), CD8+ Tem, and CD8+ central memory (Tcm) cells than naïve T-cell populations (Tn). In P2, however, both CD4+ Tem and CD8+ Tem and Tcm had robust pSTAT1 responses to IL-6. **(Figure 5b, c)**. We next used mass cytometry to compare the kinetics of IL-6-induced STAT phosphorylation in both patients with *STAT1* GOF **(Figure 5d, Supplementary Figure 4)**. Strikingly, both CD4+ and CD8+ T cells in P2 responded robustly to IL-6, reaching maximum 2.6-fold p-STAT1 above baseline at 15 minutes after induction. This response was maintained for over 90 minutes, with a significant response maintained to at least 120 minutes after stimulation. In comparison, the STAT1 response in P1 peaked at 45 minutes post-stimulation before slowly falling to baseline after the 120-minute time course. IL-6 thus can thus lead to robust and sustained activation of pSTAT1 that is in excess of that induced by IFNγ and is exacerbated in *STAT1* GOF patients and may contribute to disease pathogenesis.

## Discussion

We have identified profound and specific impacts on the immune systems of patients with *STAT1* GOF mutations through deep immune profiling that included enhanced and altered T cell activation and an overall shift in the balance of memory and Th17 T cells. The findings here highlight a unique immune phenotype and pattern of cytokine response that may play a role in T cell differentiation and memory in *STAT1* GOF patients. Cell surface and metabolic markers suggested that both patients with *STAT1* GOF mutations had a more activated phenotype and at least in the Tem cells, an exhausted T cell phenotype.

We also demonstrated a metabolic phenotype in the two patients with *STAT1* GOF. The Glut family of facultative glucose transporters are a key component of metabolic control, affecting T-cell differentiation and activation [25]. Glut1 plays a key role for effector T cells, but is dispensable for Treg cells, and may instead diminish their function [25]. The increased expression of Glut1 in *STAT1* GOF Tregs may indicate a more effector and less suppressive phenotype, and when combined with the lower percentage of Tregs in our patients, may be responsible for the increased risk of autoimmune disease. Conversely, CPT1a protein is responsible for catalyzing the rate-limiting step of fatty-acid oxidation (FAO) pathway. This pathway plays an important role in cell proliferation, and suppression of apoptosis. Our two patients had an overall pattern of increased Glut1 and CTP1a expression across immune cells, indicating an activated glycolytic and FAO metabolic program like what was recently described for early activated T cells [9] despite that neither patient had clinical evidence of infection or inflammation.

The Jak-STAT pathway is responsible for reacting to extracellular signals necessary for T cell proliferation and function [26, 27]. IL-6 has been described to activate STAT1 and STAT3 [27], with STAT3 activation leading to T-cell recruitment and survival, as well as maintenance of activated T cells in inflammatory tissues [28–31]. IL-6 activation of STAT1 is more regulatory and determines transcriptional output of STAT3 [32–35]. These activities suggest that IL-6 affects CD4+ T cell memory [27, 36–39]. While IL-6 signaling is not required for the generation or maintenance of memory T cells [36, 40], it has been demonstrated to promote the effector characteristics of CD4+ Tem [27]. Phosphorylation of STAT1 in response to IL-6 is also diminished in activated and memory CD4+ T cells [27]. In a murine model of the role of STAT proteins in activated T cells [27], strong IL-6 induction of p-STAT1 was observed in naïve CD4+ T cells but was less in CD4+ Tcm and Tem cells. The regulation of IL-6 induced STAT1 phosphorylation in activated cells has been proposed to be limited by the tyrosine phosphatases PTPN2 [27]. We hypothesize that the decreased rate of p-STAT1 dephosphorylation in P2 may similarly result in elevated and prolonged pSTAT1 signaling and impaired memory formation leading to his clinical phenotype, in particular his ability to not develop disease with initial infection but pathology with recurrent HSV and VZV.

*STAT1* GOF is associated with a heterogeneous range of clinical presentations and findings. The development of HLH is an infrequent manifestation of *STAT1* GOF mutations [41]. The mechanism of HLH development in *STAT1* GOF is unknown, but possible suggested pathways include impaired NK cell function as seen in familial HLH [42] and abnormal IFNγ signaling given the dominant role of this cytokine in driving immune dysregulation in patients with impaired cytoxicity [43]. Using our stimulation panel, we were able to demonstrate that IL-6 leads to a differing pattern of p-STAT1 response than IFNγ stimulation in the patients with *STAT1* GOF mutations than previously described [43]. While recognizing our small sample size, given the degree of signal intensity in response to IL-6, follow up assays are warranted to determine if IL-6 is a potential additional target for control of the consequences of increased p-STAT1 in this patient cohort.

The role of IL-6 may become imperative when discussing therapeutic interventions for this patient population. Treatment recommendations for patients with *STAT1* GOF are not well established and include ruxolitinib and curative bone marrow transplant. NK cell dysfunction and/or deficiency is well described in *STAT1* GOF patients [42] and is at least partially responsive to ruxolitinib [3]. Long-term outcomes with JAK inhibition are lacking and transplant survival rates to date have been very poor compared to other immune diseases including familial HLH [41]. With identification of IL-6 as a potential pathway to drive *STAT1* GOF activity, further studies will need to be performed to determine if blockade can result in better immune modulation, particularly those patients with HLH.

*STAT1* GOF mutations are rare and a limitation to our study is reliance on a two-patient cohort. As is the case when using patient samples, there are limitations to assays that can be performed and samples are reflective of the state of the patient at the time of the blood drawn, including environmental exposures, and other external factors. As such, we recognize that the phenotype of these patients could be affected by both cell intrinsic and extrinsic processes. Notably, across all the stimulation assays P1 had a slightly diminished overall response in p-STAT1 compared to P2 and healthy individuals. These data suggest a general defect or potential repressive mechanism for cytokine signaling in this patient. While prior immune modulation with corticosteroids may have affected his cellular response, he had not received corticosteroids for 6 months prior to study samples being obtained. One possible explanation is that P1 has an intrinsic cellular defect in the ability to respond to certain cytokine stimulation. It has been demonstrated that repetitive IFNγ stimulation in *STAT1* GOF results in a blunted response [20]. If a similar mechanism occurred in P1, he may have developed an improper IFNγ dependent immune challenge to histoplasmosis resulting in disseminated infection and would also explain his repressed *in vitro* p-STAT1 signaling to IFNγ in our experiment.

Our findings based on two patients with *STAT1* GOF mutations in the coiled-coil domain demonstrate that despite differing clinical presentations including age at first disease manifestation, many immune phenotypic features were shared by both patients. These shared phenotypes may be fundamental to *STAT1* GOF pathology. Additionally, we demonstrate an activated phenotype and metabolism across T-cell populations that could provide insight into mechanisms of disease, and why some patients have differing clinical phenotypes. With identification of IL-6 signaling playing a potential role in T-cell maturation, further studies will need to be performed to determine if IL-6 modulation could be used as a treatment modality. Finally, deep immunophenotyping of patients with inborn errors of immunity (IEIs) such as *STAT1* GOF can point to new disease mechanisms of human immunity and inflammation and lead to improved knowledge of cytokines and cellular signaling, such as the IL-6 pathway, in normal and pathogenic hematopoiesis and cellular maturation.

## Methods

### Donor PBMC Collection

Peripheral blood mononuclear cells (PBMC) were collected from healthy volunteers with written informed consent under IRB protocols #131311 and #191562. The patients with *STAT1* GOF were collected with written informed consent under IRB protocol #182228. Samples were de-identified prior to processing, and no other information was obtained from healthy individuals outside of a brief health survey.

### Tissue Collection and Processing

Blood was collected by venipuncture into heparinized collection tubes (Becton Dickinson; 100 mL/donor). Whole blood was diluted 1:4 with PBS before being overlaid onto a Ficoll-Paque Plus density gradient (GE Lifesciences) Blood was then centrifuged at 400g for 30 minutes without braking. Buffy coats were isolated, washed with PBS, and centrifuged at 500g for 10 minutes. Cell pellets were then resuspended in ACK lysis buffer for 5 minutes, washed, and cryopreserved at 1X10^7^ cells/mL at −80°C in 10% DMSO in FBS.

### Metal-isotope Tagged Antibodies

All antibodies used for mass cytometry analysis are listed in **Supplementary Table 2**. Pre-conjugated antibodies to metal isotopes were purchased from Fluidigm or from commercial suppliers in purified form and conjugated in house using the Maxpar X8 chelating polymer kit (Fluidigm) according to the manufacturer’s instructions procedures.

### Cell Preparation and Mass Cytometry Acquisition

Cryopreserved samples were rapidly thawed in a 37°C water bath and resuspended in complete RPMI supplemented with 10% FBS and 50 units/mL of penicillin–streptomycin (Thermo Scientific HyClone). Cell suspensions were processed and stained as previously described [14, 44, 45]. Briefly, cells were washed once with serum free RPMI and subsequently stained with ^103^Rh Cell-ID Intercalator (Fluidigm) at a final concentration of 1 uM for 5 minutes at room temperature. Staining was quenched with complete RPMI before washing with PBS 1% BSA. Cells were resuspended in PBS/BSA and added to the appropriate antibody cocktail of cellsurface staining antibodies (**Supplementary Table 2)** and incubated at room temperature for 30 minutes. Samples were washed in 1% PBS/BSA before fixation in 2% paraformaldehyde for 10 minutes at room temperature. Cells were again washed in PBS and fixed in ice cold methanol with gentle vortexing before storage at −20°C. On the day of data collection, samples stored at −20°C were washed in PBS/BSA and resuspended in an antibody cocktail of intracellular stains for 30 minutes. Iridium Cell-ID Intercalator was added at a final concentration of 125 nM and incubated at room temperature for at least 30 minutes. Cells were then washed and resuspended in ultrapure deionized water, mixed with 10% EQ Four Element Calibration Beads (Fluidigm) and filtered through a 40 uM FACS filter tube before data collection. Data were collected on a Helios CyTOF 3.0 (Fluidigm). Quality control and tuning processes were performed following the guidelines for the daily instrument operation. Data were collected as FCS files.

### Cytokine Stimulation and Phospho-specific Cytometry

Phospho-specific mass cytometry was performed as previously described [14]. Cryopreserved samples were thawed in a water bath as above. Cell pellets were resuspended in complete RPMI (10% FBS, penicillin-streptomycin) and rested at 37°C for 15 minutes. Cell suspensions were then washed in PBS and stained in 1 μM rhodium Cell-ID intercalator in PBS for 5 minutes. Cells were again washed in PBS/BSA and aliquoted equally into cytokine stimulation solutions. Briefly, these conditions included PBS (unstimulated), recombinant human interferon alpha (20 ng/mL), recombinant human interferon gamma (20 ng/mL) recombinant human interleukin 6 (20 ng/mL), recombinant human interleukin 10 (20ng/mL) or hydrogen peroxide (10uM). Stimulation conditions were allowed to proceed for indicated times before immediate fixation in 2% PFA in order to halt phospho-protein dissociation. Samples were then washed in PBS/BSA, stained with a cocktail of cell-surface antibodies and fixed in ice cold methanol as above. On the day of collection, samples were stained with a cocktail of intracellular antibodies. Iridium intercalation, resuspension in EQ calibration beads, and sample collection proceeded as above.

Fluorescence cytometry was performed using a similar, previously published fluorescence cell barcoding protocol [46]. Cells were then analyzed using a 5-laser LSRII fluorescence cytometer (BD Biosciences).

### Data Preprocessing

Raw mass cytometry files were normalized using the MATLAB Bead Normalization tool [47]. Files were then uploaded to the cloud-based analysis platform Cytobank. Before automated high-dimensional data analysis, the mass cytometry data were transformed with a cofactor of 5 using an inverse hyperbolic sine (arcsinh) function. Cell doublets were first excluded using Gaussian parameters (Center, Offset, Width, Residual) as reported [14]. Intact cells were gated based on DNA content (^191^Ir and ^193^Ir). Dead cells were excluded based on rhodium intercalation. Immune subsets were then manually gated using biaxial gating strategies.

Fluorescently barcoded samples were debarcoded and assigned to individual wells on a 48-well plate using the DebarcodeR algorithm[48].

### Dimensionality Reduction

Analysis by t-distributed stochastic neighbor embedding (t-SNE) was performed using the Cytobank platform. For each respective panel, 800-31,000 total cells of interest (CD45+CD3+ T cells, CD45+CD19+ B cells, or CD45+CD3-CD19-CD56-myeloid cells) were pregated before analysis. Dimensionality reduction was performed using all markers within each panel. Metabolic and phospho-protein markers were excluded to assess individual marker expression. Each t-SNE map was generated using a perplexity of 30, theta of 0.5 and 10,000 iterations.

### Statistical Analysis

All statistical analysis was performed in Cytobank, R version 3.6.1, or Graphpad Prism version 8.4.3. Where appropriate, median intensity values were transformed using the arcsinh scale normalized to the minimum intensity value across all markers. Patient values outside of the IQR of five pooled healthy donor samples were considered significant.

### Study Approval

Written informed consent was obtained from all participants prior to participation, and this study was conducted in accordance with the Declaration of Helsinki, approved by the IRB at Vanderbilt University Medical Center.

## Author contributions

S.K., T.B., J.A.C., J.M.I., and J.C.R. designed the data science study. S.K. was first co-author given the translational questions posed as she worked to phenotype these patients clinically, and then asked the questions that were then answered via the research platform. M.H., D.D., S.S. performed experimental work. S.K., T.B. performed data analysis, developed figures, and wrote the first draft of the manuscript. S.K., T.B., J.A.C., D.E.D, J.M.I. and J.C.R. edited the manuscript. S.K., D.E.D, Y.E.K, and J.A.C. compiled patient data. All authors read and approved the manuscript.

## Acknowledgements

We would like to thank our patients and their families for their participation in this work, and their willingness to contribute to our understanding of inborn errors of immunity. We also thank members of the Rathmell and Irish labs as well as members of the Human Immunology Discovery Initiative and Summer Brown for administrative and IRB support.

## Compliance with Ethical Standards

Informed consent was obtained from all participants, and this study was conducted in accordance with the Declaration of Helsinki, approved by the IRB at Vanderbilt University Medical Center.

## Supplementary Figures

**Supplementary Figure 1.**
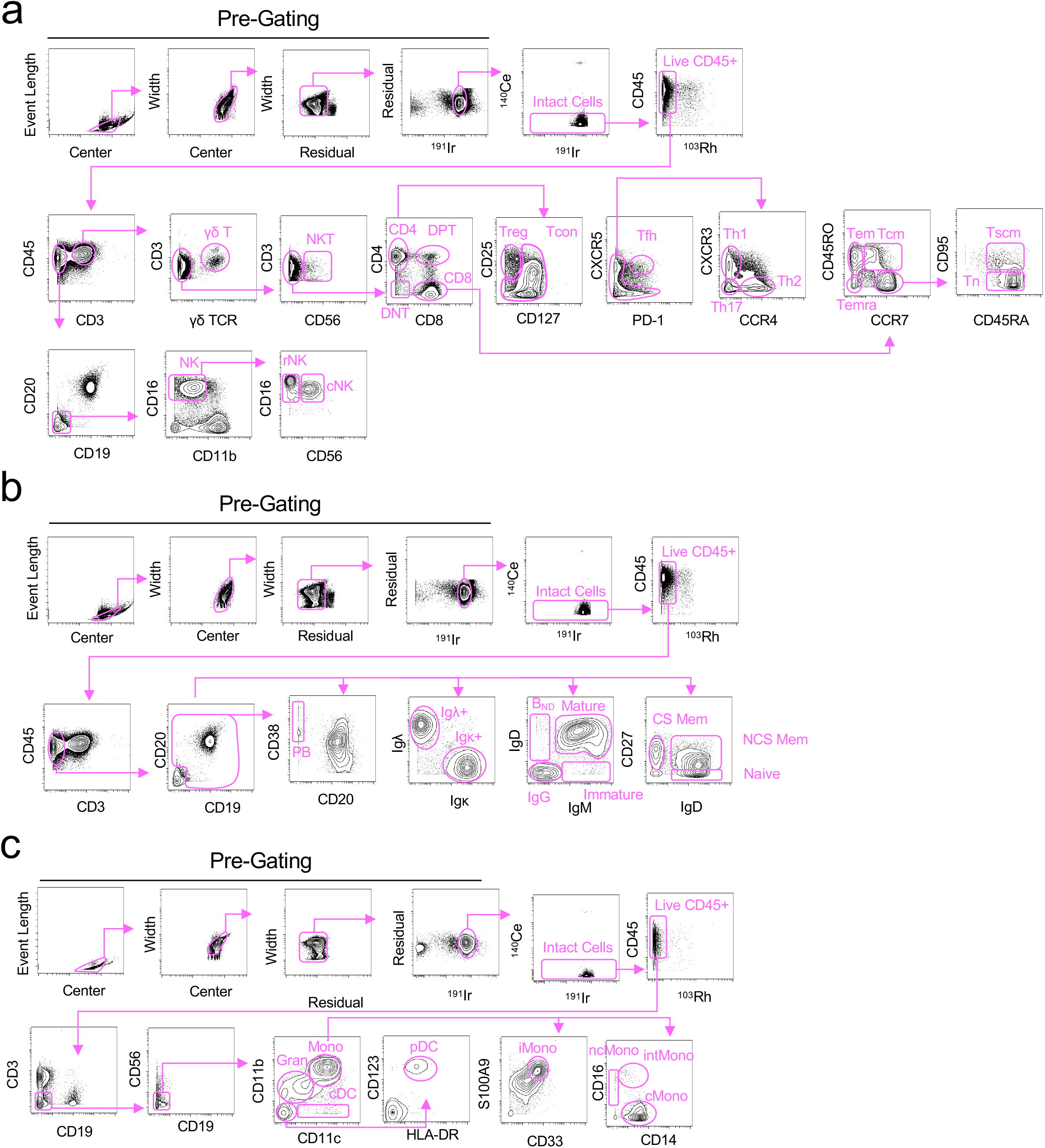
Immune Cell Gating Schema. Representative gating schema for expert identification of T cell subsets **(a)** B cell subsets (**b**), or myeloid subsets (**c**). All populations were pre-gated using Gaussian parameters before expert gating on individual cell populations. Normalization beads were gated out using the ^140^Ce channel. Live, intact cells were gated using the ^103^Rh ^191^Ir DNA intercalators respectively. γδ T= gamma delta T cell. NKT= Natural Killer T cell. DNT= Double negative T cell. DPT= Double Positive T cell. Treg= regulatory T cell. Tcon= Conventional helper T cell. Tfh= Follicular Helper T cell. Th1= T helper 1. Th2= T helper 2. Th17= T helper 17. Tem= Effector memory T cell. Tcm= central memory T cell. Temra= Effector memory CD45RA+ T cell. Tn= Naïve T cell. Tscm= stem cell memory T cell. NK= Natural Killer cell. rNK= regulatory NK cell. cNK= cytotoxic NK cell. PB= plasmablast. Igλ= immunoglobulin lambda. Igκ= immunoglobulin kappa. BND= naïve IgD+ B cell. CS Mem= class switched memory B cell. NCS Mem= non-class switched memory B cell. Gran=granulocyte. Mono=monocyte. cDC=classical dendritic cell. pDC=plasmacytoid dendritic cell. iMono= inflammatory monocyte. ncMono=non-classical monocyte. intMono=intermediate monocyte. cMono=classical monocyte.

**Supplementary Figure 2.**
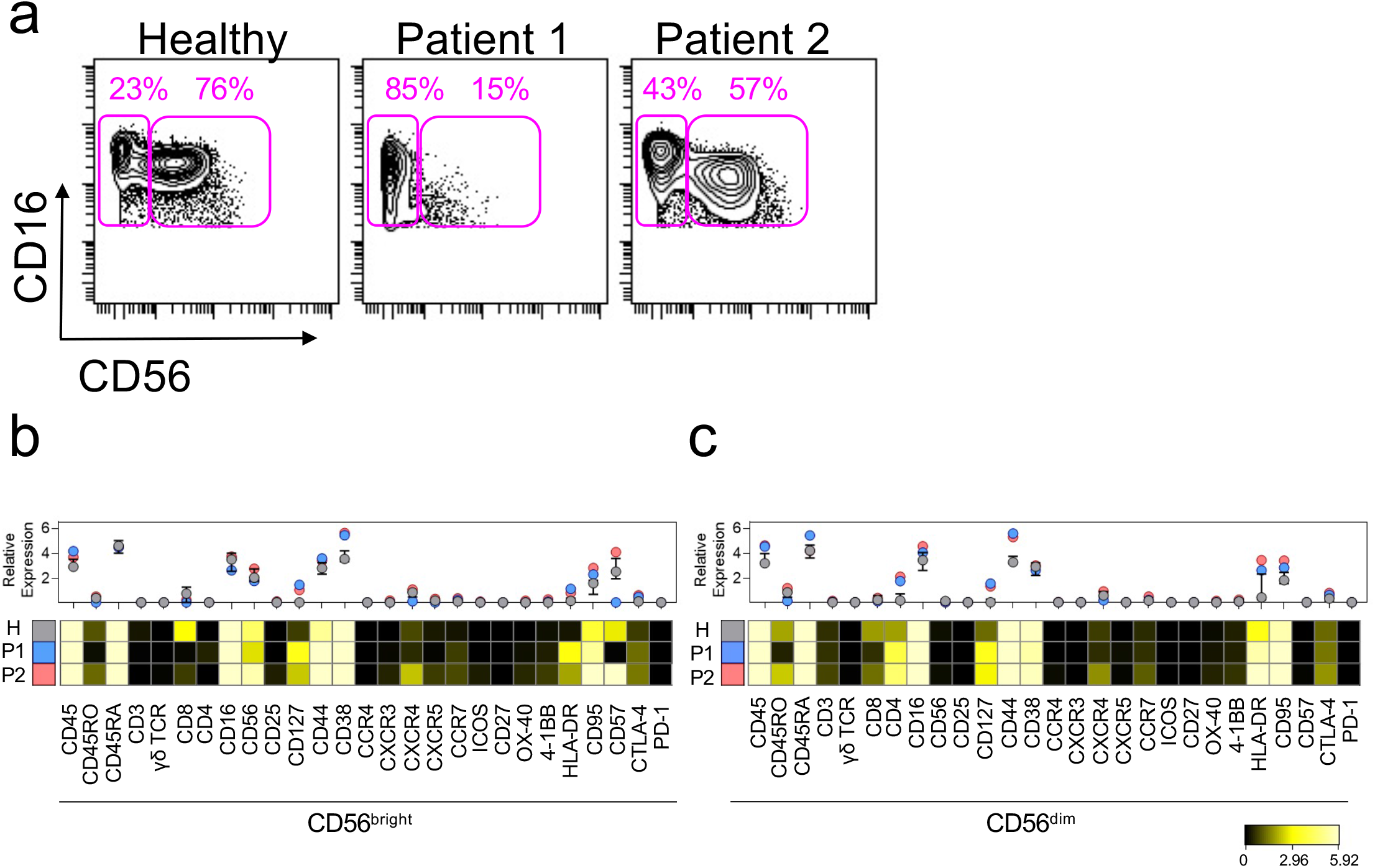
Quantification of NK cell populations in patients with *STAT1* GOF mutations. **a**. Representative biaxial plots indicating the frequencies of CD16+CD56dim regulatory NK cells and CD16+CD56bright cytotoxic NK cells in healthy donor or patient PBMC. Heat maps indicating the arcsinh transformed median values of 27 phenotypic markers expressed in manually gated CD56bright (**b**) or CD56dim (**c**) NK cell populations. Graphs above each heatmap indicate the median arcsinh fold change expression of each marker relative to the minimum median mass intensity. All statistics represent the median value. Errors bars indicate the interquartile range.

**Supplementary Figure 3.**
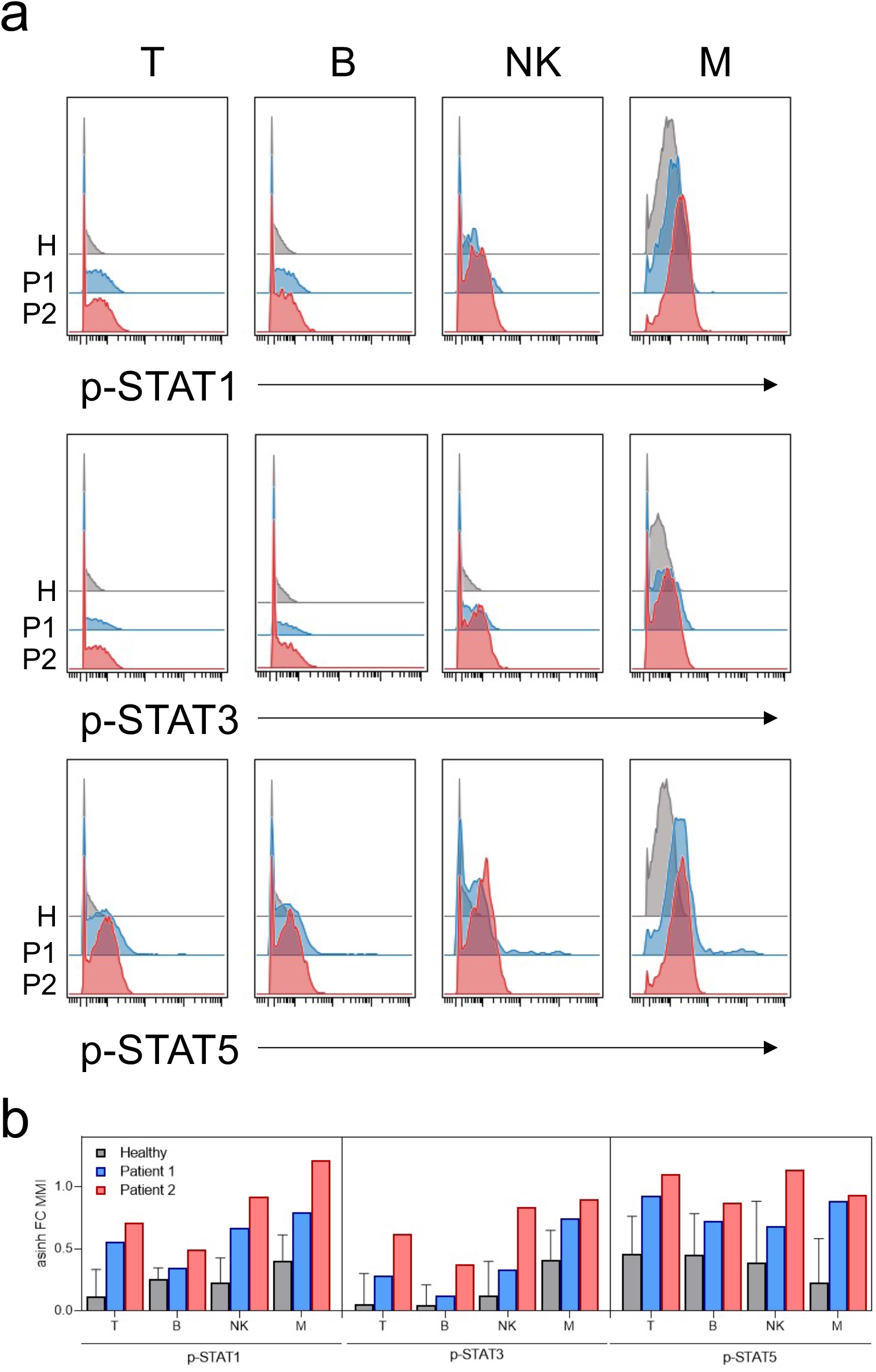
Patients with *STAT1* GOF mutations exhibit elevated basal STAT phosphorylation across multiple immune subsets. **a**. Representative histogram plots indicating the basal expression of p-STAT1 (top), p-STAT3 (middle) or p-STAT5 (bottom) within indicated manually gated immune cell populations. **b**. Quantification of the arcsinh transformed median intensity values of indicated phospho-STAT proteins normalized to the minimum expression value across all immune subsets. Bars represent median intensity values. Error bars indicated the interquartile range.

**Supplementary Figure 4.**
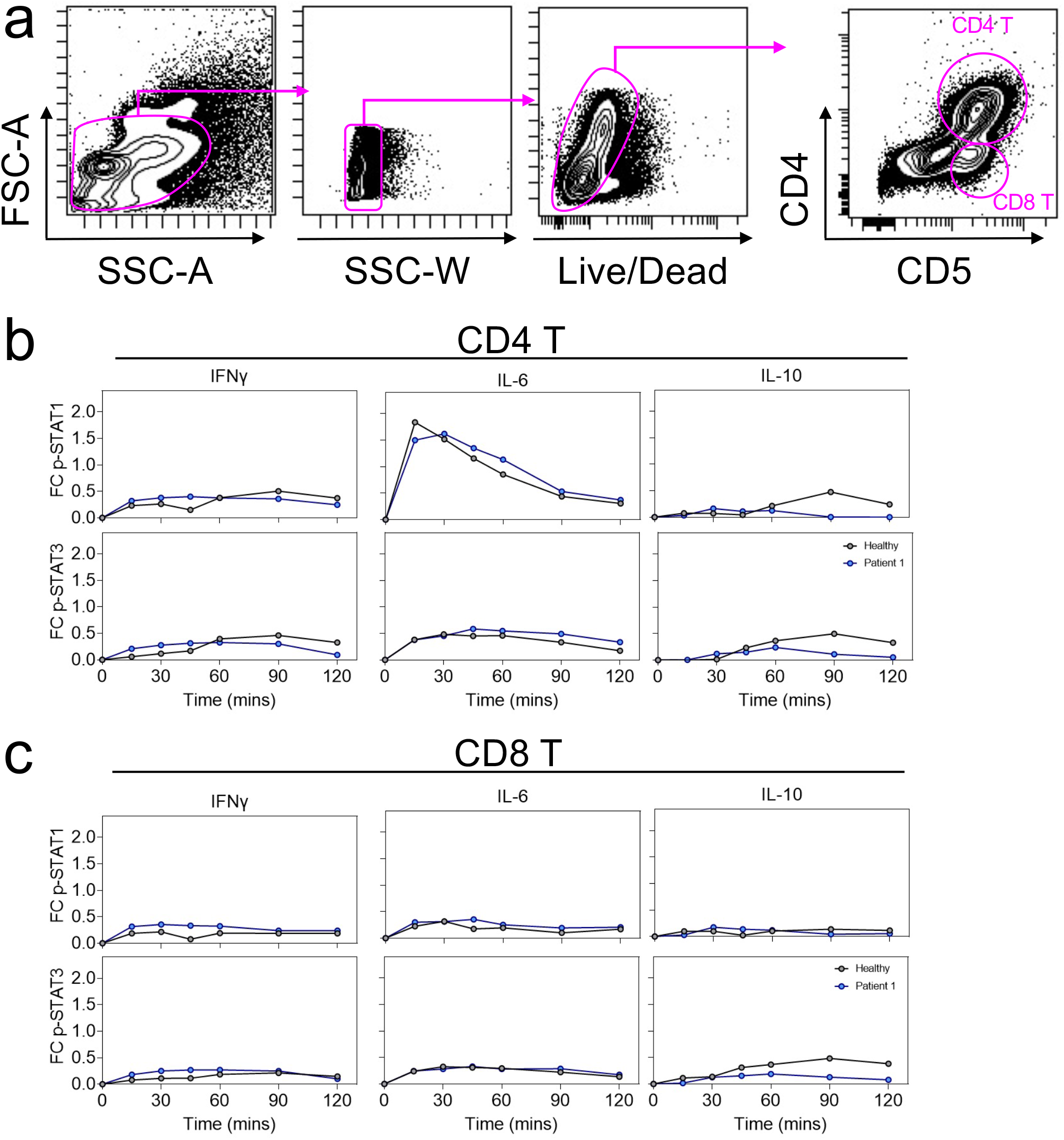
Fluorescence Flow Stimulation Assay for Patient 2 Outlines the p-STAT1 and p-STAT3 response in CD4+ and CD8+ T cells. **a**. Representative biaxial gating strategy to evaluate phospho-signaling in CD4+ and CD8+ T cells. PBMCs were cultured in either IFNγ (left), IL-6 (middle) or IL-10 (right) for the indicated length of time from 0 minutes to 120 minutes before fixation and quantification of STAT1 (top) or STAT3 phosphorylation (bottom) in CD4+ T cells **(b)** or CD8+ T cells **(c)**. The indicated p-STAT MFI was normalized and arcsinh transformed each individual’s unstimulated (0 minute) value for each T cell population.

---

**Supplementary Table 1:** Patient Characteristics

|  | PI | P2 |
| --- | --- | --- |
| Age at time of work up below | 9 months old | 9 years old |
| Sex | M | M |
| *STAT1* Mutations | c.800C>T; p.Ala267Val | c.866A>G; p.Tyr289Cys |
| Absolute Lymphocyte Count (2000-6200/mcL) | 490 | 1790 |
| CD3+ (1700-3600) | 186 #/cumm | 1342 #/cumm |
| CD4+ (1000-2800) | 108 #/cumm | 770 #/cumm |
| CD8+ (800-15000) | 69 #/cumm | 465 #/cumm |
| CD16+56+ (200-700) | 198 #/cumm | 394 #/cumm |
| CD19+ (500-1500) | 821 #/cumm | 18 #/cumm |
| T cell Proliferation to Mitogen | Normal to PHA and PWM | Normal to PHA and PWM |
| T cell Response to Candida | Decreased in CD45+, normal in CD3+ | Absent |
| T cell Response to Tetanus | Decreased | Absent |
| NK Function (chromium release assay) | Decreased | Decreased |
| CD107a degranulation | Normal | Normal |
| IgG (320-1150 mg/dL) | 412 | 1188 |
| IgM (40-150 mg/dL) | 24 | 44 |
| IgA (0-90 mg/dL) | 11 | 133 |
| Diphtheria Titers (IU/mL) | 1.7 (Protective) | 2.2 (Protective) |
| Tetanus Titers (IU/mL) | 0.8 (Protective) | 6.8 (Protective) |
| Streptococcal Pneumoniae Titers (IU/mL) | 12/13 serotypes in PCV-13 Protective | 12/13 serotypes in PCV-13 Protective |
| EBV IgM/IgG | Negative | Negative |
| CMV IgM/IgG | Negative | Negative |

**Supplementary Table 2:**
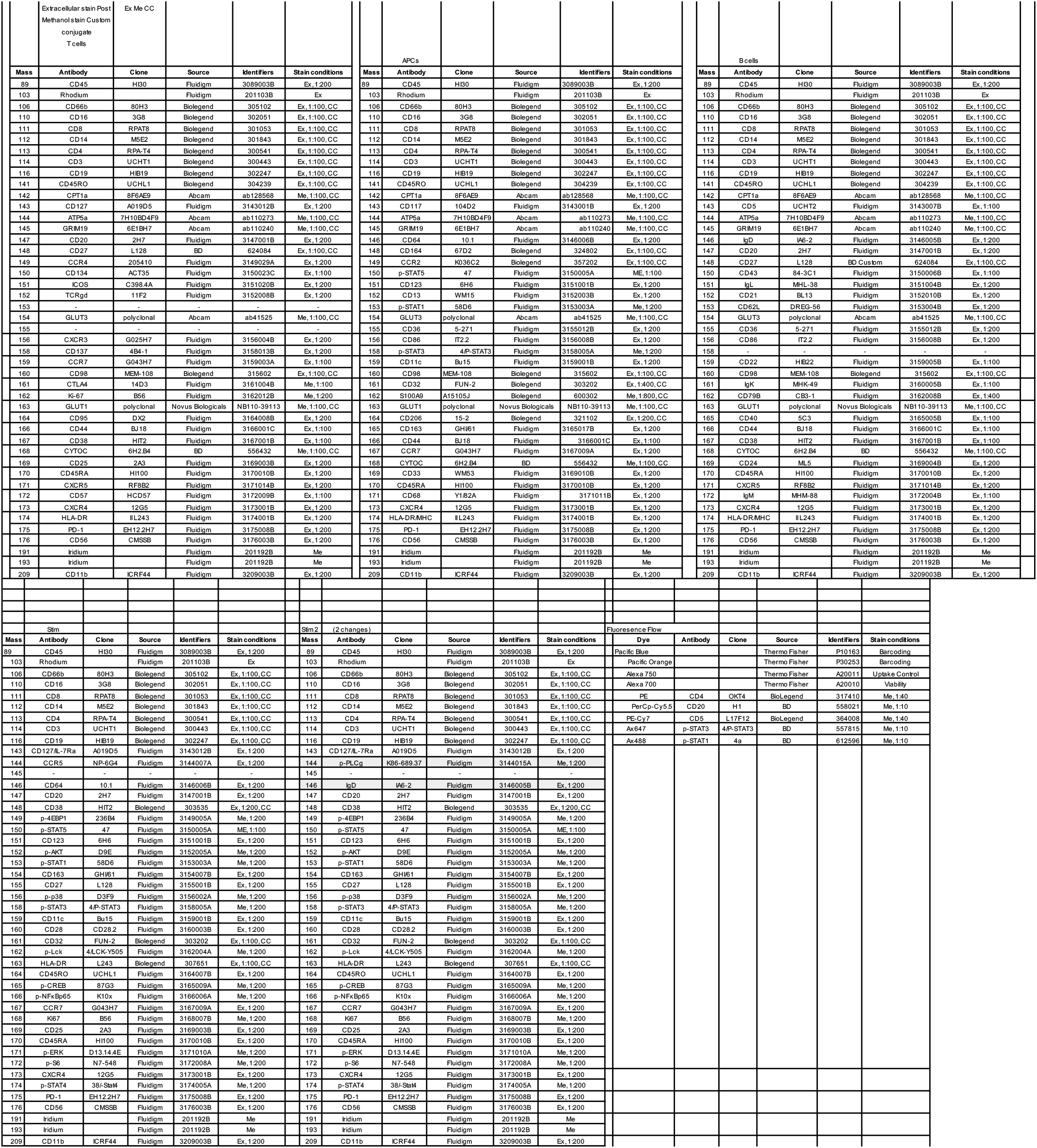
Antibody Panels

